# Fully-Automated Multicolour Structured Illumination Module for Super-resolution Microscopy

**DOI:** 10.1101/2024.07.04.601961

**Authors:** Haoran Wang, Peter T. Brown, Jessica Ullom, Douglas P. Shepherd, Rainer Heintzmann, Benedict Diederich

## Abstract

In the rapidly advancing field of biological imaging, there is a great need for high-resolution imaging techniques that are both cost-effective and accessible, for example to better observe and understand dynamics in intracellular processes. Structured illumination microscopy (SIM) is the method of choice to achieve high axial and lateral resolution in living samples due to its optical sectioning and minimal phototoxicity. However, the high cost and complexity of conventional SIM systems limit their wide application. In our work, we present an open-source, fully-automated, two-color structured illumination module that is compatible with commercially available microscope stands. The compact design, consisting of low-cost single-mode fiber-coupled lasers and a digital micromirror device (DMD), is integrated into the open-source acquisition and control software (ImSwitch) in order to realize real-time super-resolution imaging. This developed system achieves up to a 1.55-fold improvement in lateral resolution compared to conventional wide-field microscopy. To rationally design this module, we developed a model to ensure optimal DMD diffraction performance using tilt and roll pixels, thus covering a wide range of low-cost video projectors for use in coherent SIM setups. Our goal is to democratize SIM-based super-resolution microscopy by providing both comprehensive open-source documentation and a modular software framework that works with various hardware components (e.g. cameras, stages) and reconstruction algorithms. In this way, we try to upgrade as many devices as possible to the super-resolution realm.

## INTRODUCTION

Fluorescence microscopy is a fundamental technique for exploring biological processes at the micrometer scale which has revolutionized our understanding of the life sciences. However, our understanding of many important biological processes is limited by tradeoffs in fluorescence microscopy which make it challenging to image large volumes at high-resolution and high speeds. Recently, a wide variety of advanced imaging instrumentation has been developed to mitigate these tradeoffs, but these approaches often require significant technical expertise to build and operate or are prohibitively expensive for most life science groups. We address this problem for fluorescence super-resolution imaging by designing a low-cost, opensource add-on for realizing structured illumination microscopy on commercial microscope stands.

The demands, especially on optical imaging, are increasing in order to depict finer structures, larger volumes and faster dynamics, for example to better understand the interactions between virus and host or to recognize the effects on our inner immune system [1]. This often leads to an increase in the required technical expertise to build or operate a recently developed device or, considering commercial devices, an extremely high purchase price, whereby existing equipment can often not be reused.

To increase the optical resolution beyond the diffraction limit defined by Abbe, numerous research groups have developed innovative approaches over the past two decades. These include localization microscopy techniques based on photo-switching such as *d* STORM [2, 3] and PALM [4], as well as point-scanning approaches that rely on saturated molecular transitions such as STED and RESOLFT [5]. However, these approaches exploit fluorophore photophysics and thus require the selection of a specific fluorescent dye and high illumination intensity, which can lead to phototoxicity or photobleaching, all of which represent major hurdles for live cell imaging [6].

Structured illumination microscopy (SIM [7, 8]), as a widefield imaging technique, typically only offers a maximum improvement in resolution by a factor of 2, but only requires few photons, yields a compact raw data size, and is thus ideally suited for imaging living cells to better understand dynamic cellular processes.

SIM achieves its high resolution by illuminating fluorescent samples with a series of very fine periodic pattern of very high contrast and detecting images at various pattern positions for computationally reconstruction. This structured illumination down-modulates high spatial frequencies of the object into the detection passband of the microscope and the computational reconstruction unmixes several overlapping down-modulated object components and places each of them at the correct location and phase in the final superresolution image. Coherent illumination has the great advantage of providing near 100 % pattern contrast up to the highest frequencies supported by the illumination system. Although SIM has the same theoretical passband limit as a confocal microscope [9], the signal-to-noise ratio, especially at high spatial frequencies, is significantly better due to the very high contrast of the high frequency illumination. Coherent illumination is therefore a key component of SIM systems that aim for high resolution rather than optical sectioning. The main advantage of incoherent SIM systems such as the commercially available Zeiss Apotome is an improved sectioning capability similar to that of confocal microscopy.

Since its introduction [7, 8, 10], many have refined SIM, e.g. by replacing the illumination patterns of the classical two-beam interference and the resulting sinusoidal illumination in the sample plane with two-dimensional [11], speckle-like [12] or hexagonal illumination patterns [13], in order to simplify the setup or to achieve faster data acquisition [14]. Other efforts optimized the image processing algorithms e.g. using deep learning [15] or GPU-assisted reconstruction algorithms to realize real-time SIM [16]. Recently integrated solutions realize SIM illumination by means of photonic chips [17], metamaterials [18] or optoacoustic Bragg cells [19]. There are several ways to implement even classical two-beam SIM, such as using a Michelson interferometer [20] or a fiber array [21]. Besides the use of classical diffraction grating in a conjugate image plane, the use of a spatial light modulator allows a flexible and accurate, non-mechanical manipulation of the phases and diffraction orders. Liquid crystal on CMOS (LCoS) spatial light modulators (SLM’s) [22, 23] represent relatively straightforward to implement technologies, but are expensive and slow. Alternative approaches, such as the recently introduced method using galvo scanners [24, 25] to manipulate the phase and rotation in the BFP, require a high effort in aligning the optical components. Digital micromirror devices (DMD’s) on the other hand are a relatively cheap and fast alternative, but are amplitude rather than phase modulators, so they are less efficient in some configurations. However, DMD-SIM systems are more challenging to design and align due to complex diffraction effects caused by the tilt of the DMD micromirrors [26–28].

Those who do not have the necessary budget of several hundred thousand euros for the procurement of a commercial grade SIM instrument are therefore often confronted with high complexity in the realization of a SIM microscope. Since for SIM illumination, the optical path can be fairly complex, the possibility of building a super-resolution microscope is limited for non-specialists in optics. Established biological research groups can often draw from a selection of microscopes in their department or the local core facility, though few can keep up with current development in modern microscopy methods, which often means that better resolution requires the purchase of a new microscope and a corresponding budget.

The field of “Open Microscopy” [29] tries to solve this dilemma by providing detailed blueprints for replication verified by peer review and by using frugal science to keep the price low. With our recently introduced SIM module for the UC2 system [30, 31], we have presented a monolithic module that complements the cube-based 3D printed microscope with structured illumination imaging capability. However, the system is aimed more at education with a relatively small DMD display. In this work, we build on these principles to design an open-source SIM add-on for commercial microscope stands, which is suitable for addressing scientific questions.

Our work is designed to complement significant advancements in the field, such as the OpenSIM modules discussed in [23], by leveraging coherent illumination to achieve superior modulation contrast. Unlike the multi-color-based illumination of OpenSIM, which enhances existing commercial microscopes, our method not only offers improved image quality but also integrates open-source software. This approach makes it more accessible and costeffective, addressing common concerns with proprietary systems and high costs associated with other SIM technologies. Similarly, with our recently introduced (coherent) mcSIM framework [27], we present a fully open-source system that includes construction plans, CAD tools and control/reconstruction. However, it requires a significant budget and optical knowledge to replicate the setup, as high-class optical components are used and the setup of a complete microscope on the optical table is necessary, making replication for less-skilled people difficult.

To foster inexpensive access to SIM imaging beyond previous approaches, we present a fully open-sourced, dual-color (488/635nm) structured illumination microscopy add-on module that can be easily adapted to commercially available fluorescence microscopes. The add-on includes both hardware and software components and is supported by extensive documentation (including parts lists and setup videos). On the hardware side, we present an open-source illumination engine based around a DMD. With this system, we achieve superresolution enhancements of up to 1.5 times to the Abbe limit. This resolution enhancement could be further increased by using more aggressive SIM patterns.

We provide a SIM acquisition and reconstruction plugin for the Python-based open-source microscopy control software ImSwitch [32], enabling live stream SIM image observation and further seamless integration in automated microscopy workflows.

Our add-on is based on low-cost components, which significantly reduces the overall cost so that the final build can be done for around three thousand euros. This, together with the reduced complexity of the build process, increases the reproducibility of our approach. The concept of open-source makes this project accessible to all in need of advanced imaging systems, rendering this cutting-edge technology affordable for a broad range of research laboratories.

## METHODS

### A. Multicolor DMD alignment

In this work, we rely on a DMD to generate structured illumination patterns due its low cost, large spectral acceptance range, fast speed, and reproducible quality. As discussed above, DMD’s are beam shaping devices which are commonly used in commercial video projectors and have recently been applied to a wide variety of biomedical applications, including SIM [26–28, 33, 34]. However, DMD’s are primarily designed for use with incoherent light, and working with coherent light introduces several challenges. Most significantly, the complex DMD diffraction hampers their use in multicolor laser systems as compared with LCoS SLM’s. Additionally, DMD’s can introduce aberrations into the illumination system both due to the mirror arrays not being optically flat and also the presence of a thick glass window which is a few millimeters away from the vacuum-sealed micromirrors.

A DMD consists of a rectangular array of micromirror “pixels” arranged on a square lattice in the DMD backplane. Each mirror is mounted on a micromechanical swivel which is controlled by electrodes that can dynamically position the mirror in one of two orientations, which we refer to as the “+” or “-” orientation. The mirrors can be independently controlled to display an arbitrary binary pattern. Brightness in video-display applications is typically controlled by temporal multiplexing. Due to the tilt of the micromirrors, DMD’s are essentially blazed diffraction gratings, which are designed to optimize diffraction efficiency into a non-zero diffraction order.

A coherent plane wave incident on a DMD with mirrors all in the + state will be diffracted into many different orders (Fig. 2A) due to the periodic mirror structure. The relative intensity diffracted into the various orders is modulated by an envelope function, the blaze envelope, which has its maximum at the direction of specular reflection from the mirrors. Achieving efficient diffraction requires one of the diffraction orders to be aligned with the center of the blaze envelope, in which case we say the system satisfies the blaze condition. When the DMD displays a pattern, using mirrors in both the + and states, the pattern structure generates additional subdiffraction orders about the main diffraction orders (Fig. 2B). If the primary diffraction order does not meet the blaze condition, the relative strength of positive and negative suborders will be distorted. This distortion is particularly deleterious in SIM because unequal suborders results in reduced SIM modulation contrast, which rapidly degrades the recoverably superresolution information.

Multicolor DMD operation is particularly challenging because the output angle of the diffraction orders are wavelength dependent, but the blaze envelope direction is not. As such, multicolor operation requires either carefully tuning the illumination wavelength [35], using different input directions for different wavelengths [27], or accepting reduced SIM modulation contrast for certain wavelengths [34]. This problem is unique to DMD’s, as LCoS SLM’s are usually operated using the zeroth diffraction order, which does not exhibit chromatic dispersion.

Recent work has surmounted these challenges to realize multicolor coherent DMD-SIM using up to 3-colors [27, 35]. This work fully characterized the possible multicolor solutions which satisfy the blaze condition, but using a model specialized to DMD’s with corner illumination pixels (CIP’s). CIP’s have a rotation axis at a 45^°^ angle to the mirror lattice but in the same plane, and rotate symmetrically about this axis to orient the mirror normals ±12^°^ away from the DMD backplane normal to reach their + and - states respectively (section S6 C). However, in this work we use the widely-available and cost-effective DMD evaluation module DLP4710EVM-G2 (Texas Instruments, USA, TX) which features tilt and roll pixels (TRP’s) [36]. TRP’s use a composite rotation which effectively rotates the mirrors ≈17^°^ about the *x* and *y* mirror lattice axes in the + and - orientations respectively (section S6 D).

To rationally design our multicolor coherent DMD-SIM module, we extend the CIP diffraction model of [27] to DMD pixels with an arbitrary rotation axis, which include TRP’s as a special case. We leverage this model to select a geometry which enables 2-color SIM using 488 nm and 635 nm excitation.

To develop this model, suppose our DMD mirror lattice has pitch *d* and the mirrors rotate about an arbitrary axis 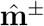 through angle γ^±^ as described by a rotation matrix 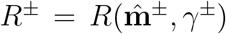 bto reach the ± orientations (section S6 B). Define a coordinate system with unit vectors ê_*x*_ and ê_*y*_ along the principal axes of the mirror lattice in the backplane of the DMD chip and ê_*z*_ normal to this plane and pointing away from the DMD. The DMD is illuminated by a coherent plane wave with incident direction described by unit vector â, and we will consider light diffracted into output direction unit vector 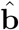 (Fig. 2b). By construction, *a*_*z*_ and *b*_*z*_ are along the ‘ê_*z*_ directions respectively,

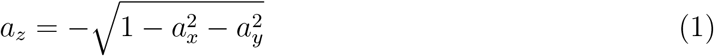

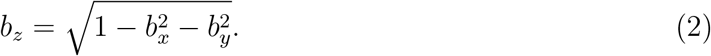

The incoming and outgoing directions must satisfy the diffraction condition of the mirror lattice,

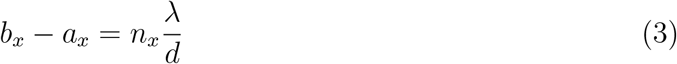

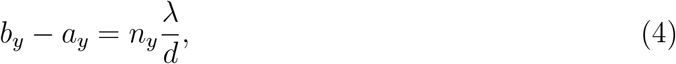

where *n*_*x*_ and *n*_*y*_ are integers indexing the diffraction orders.

To achieve high diffraction efficiency, the incoming and outgoing directions should also satisfy the blaze condition. To simplify our analysis of this condition, we introduce a new coordinate system, defined by unit vectors 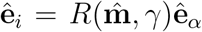 for *i* = 1, 2, 3 and α = *x, y, z*. Since ê_3_ points along the micromirror normal, the blaze condition in these coordinates is

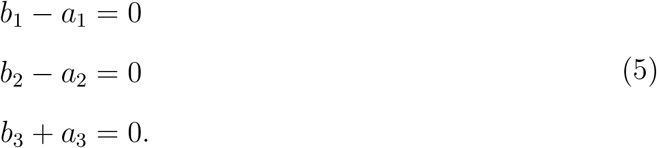

Combining the blaze and diffraction conditions, we have six unknowns, the components of â and 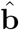, and seven equations. The system is over-determined and does not have any exact solutions in the general case. However, for specific choices of the mirror rotation matrix the above system of equations can have an exact solution. To understand when exact solutions exist, we rewrite eqs. 3 and 4 in the mirror basis and substitute eqs. 5 to find

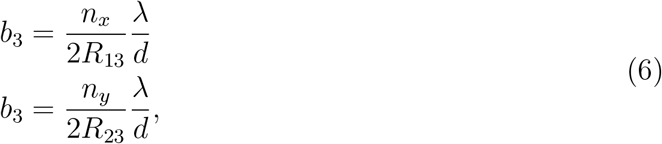

where *R*_*ij*_ are the matrix elements of the mirror rotation matrix. Eqs. 6 can only be satisfied if the ratio of *R*_13_ and *R*_23_ is a rational number. For example, for CIP’s *R*_23_ = -*R*_23_ implying exact solutions exist for diffraction orders *n*_*x*_ = -*n*_*y*_ (section S6 C).

In the general case, we can alternatively view this as an optimization problem where we regard the diffraction equations as constraints and the blaze condition violation as a function we would like to minimize. To this end, we define a cost function

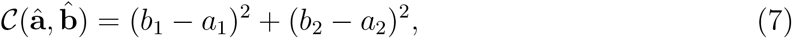

which characterizes how much input-output direction pair â and 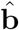 violate the blaze condition. This cost function is not unique, as we could include additional terms penalizing deviations in *b*_3_ + *a*_3_. But as we will see, this choice is expedient because it is easy to solve, and it reproduces the exact solutions if they exist.

We minimize the cost function subject to our constraints using the method of Lagrange multipliers, which supplies us with a new set of equations to replace the blaze condition,

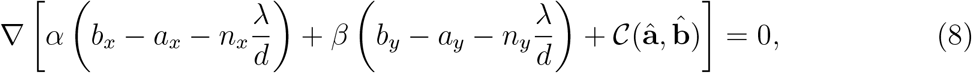

Where α and β are Lagrange multipliers, and the gradient is taken with respect to *a*_*x*_, *a*_*y*_, *b*_*x*_, and *b*_*y*_. The details of solving this system are provided in (section S6 A).

For TRP’s in the - state, if we divide equations 6 we find 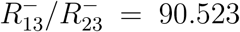 (section S6 D), where *R*^-^ is the rotation matrix describing the mirror orientation for the - state mirrors. This implies there are no exact solutions, because even if the ratio is rational, the large diffraction orders required to satisfy it are not physical. We can search for approximate solutions for diffraction orders where this ratio is large. In particular,

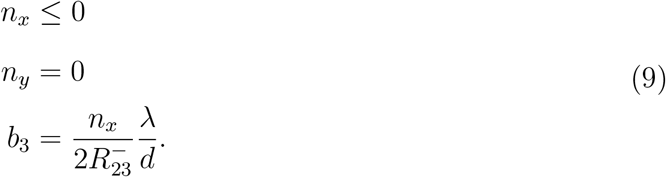

These solutions would be exact, except for the small component of the rotation axis along the ê_*z*_ direction. The situation for the + state is identical under 90^°^ rotational symmetry.

We solve this system of equations for the TRP pixels with both 488 nm and 635 nm to determine the optimal input and output directions and find two-color compromise solutions (Fig. 2e). Since each choice of mirror orientation and diffraction order produces a different solution, we solve for both the + and - orientations and a sequence of low diffraction orders matching the form determined in eq. 9, in particular (*n*_*x*_, *n*_*y*_) = (*n*, 0) for the - mirrors, and (0, *n*) for the + mirrors, where *n* = -1, …, -6. For most diffraction orders, we find that the solution directions are not unique, but form a one-parameter family. We display these solutions as curves in output unit vector space, parameterized by *b*_*x*_ and *b*_*y*_ (Fig. 2E). The corresponding input directions for any point on these curves can be determined from the diffraction conditions, eqs. 3 and 4. For 488 nm illumination, the innermost closed curve corresponds to the (0, -6) order, and the larger curves correspond to progressively decreasing orders. For the 635 nm illumination, the solution for the (0, -6) and (0, -5) orders are single points, and the innermost closed curve corresponds to the (0, -4) order. The larger curves again correspond to progressively decreasing diffraction orders.

While the solutions displayed in Fig. 2e) are all optimal in the sense of eq. 7, they do not have identical blaze condition violation angles, Δ_*b*_ because eq. 7 does not consider the ê_3_ direction. Furthermore, solutions for different diffraction orders may have significantly different blaze angle violations. However, for the TRP pixels we find the blaze condition violations angles are typically ≤0.5^°^, and the magnitude is shown using the color scale in Fig. 2E.

From the simulations, we conclude that the TRP - mirror state supports nearly-blazed 2- color solutions for the 488 nm (−5, 0) order and the 635 nm (−4, 0) order which nearly overlap due to the fact 635*/*488 ≈ 5*/*4. While there are many possible choices of input/output directions, for experimental convenience, it is convenient to work with directions where the beams stay in the same 2D plane. For the TRP – state, this corresponds to light incident in the *xz* plane. Focusing on this plane, we have a solution with output direction that forms an angle of ≈20^°^ with the DMD backplane normal. For 488 nm the optimal solution is incident at an angle of 55.9^°^ and diffracts at an angle of 22.1^°^ with a blaze angle violation of 0.29^°^. For 635 nm the optical solution is incident at an angle of 52.9^°^ and diffracts at an angle of 19.1^°^ with a blaze angle violation of 0.3^°^. The input angle of one or both beams must be adjusted to ensure the output angles are the same, entailing some additional blaze angle violation. For example, if the 635 nm beam is aligned to the 488 nm beam, the incidence angles must be modified to 57.9^°^, and the blaze condition violation increases to 1.97^°^.

While the previous solution achieves small blaze condition violation, it requires the DMD backplane to be tilted significantly with respect to the optical axis, which can introduce unwanted effects including optical aberrations and pattern focal plane variation across the field of view. To address these issues, we also consider solutions for the 488 nm (−6, 0) order and the 635 nm (−5, 0) order. In the optimal alignment for the 488 nm (−6, 0) order, the incoming beam is incident at 38.1^°^ with blaze violation of 0.34^°^ and DMD backplane tilt of 4.3^°^. Aligning the 635 nmorder to the same output direction as the 488 nm beam results in an incidence angle of 41.6^°^ with a blaze violation of 3.4^°^. We choose to use this solution in the experiment.

Previous experiments have reported DMD mirror rotation parameters which differ somewhat from the nominal values [16, 26, 27]. To address this possibility, we infer the true mirror rotation parameters for the - state of our DMD by displaying a series of structured patterns on the DMD, recording the intensity of their various diffraction sub-orders on a camera, and fitting our model to the results [27] (section S6 E). We determine that the mirror rotation axis is ≈13^°^ different than the nominal value, and the mirror rotation angle is 17.7^°^. These differences imply typical optical blaze condition violations are on the order of 3^°^, somewhat larger than the values obtained for the nominal DMD parameters we considered above.

### B. Mechanical and Optical Design of the SIM Setup

Our goal is a compact, low-cost setup that adapts with a large variety of different microscopy bodies. The developed hardware represents a compromise between price, availability, complexity and achievable resolution enhancement. A laser-tight custom-designed DIY enclosure ensures that lenses, mirrors, and SLM have predetermined positions, streamlining the assembly process. This reduces the alignment effort to a minimum, with assembly by semi-experienced opticians being possible in about 2 hours (provided that all parts are prepared in advance), based on detailed step-by-step documentation [37]. Using different adapter plates and tube lenses respectively enables the adaptation to different microscopy bodies. To demonstrate the openness of our setup, we provide an adapter for the OpenFrame project [38], promoting open science by enhancing an existing system with an imaging modality instead of developing a new one. This increases the likelihood of implementation due to the widespread use of OpenFrames (see Fig. S6). A complete bill of materials can be found in Sec. S8).

For the light source, we employ two single-mode fiber-coupled (core 4 µm, non-polarization maintaining) diode lasers (openUC2, Jena, Germany) with operating wavelengths of 488 nm and 635 nm. Their output power is tuned via PWM signals from an ESP32 microcontroller (binary pulse width modulated, PWM, signal at *f*_duty cycle_ = 5 kHz at 10*Bit* resolution).

The optical setup geometry, as illustrated in Fig. 1, was designed with the help of the previously calculated optimal beam incidence angles for the 488 nm (−6, 0) and the 635 nm (−5, 0) orders using the - mirrors. The laser light emerging from the single-mode fibers is collimated using achromatic lenses of 50 mm focal length (AC-254-50-A M, Thorlabs). Then the two beams are combined and directed towards the DMD chip using mirrors (PF10-03- P01, Thorlabs), which are arranged to minimize the overall footprint of the assembly. The DMD displays the desired SIM patterns, and we direct light diffracted from the - mirrors towards the collection optics.

**FIG. 1.**
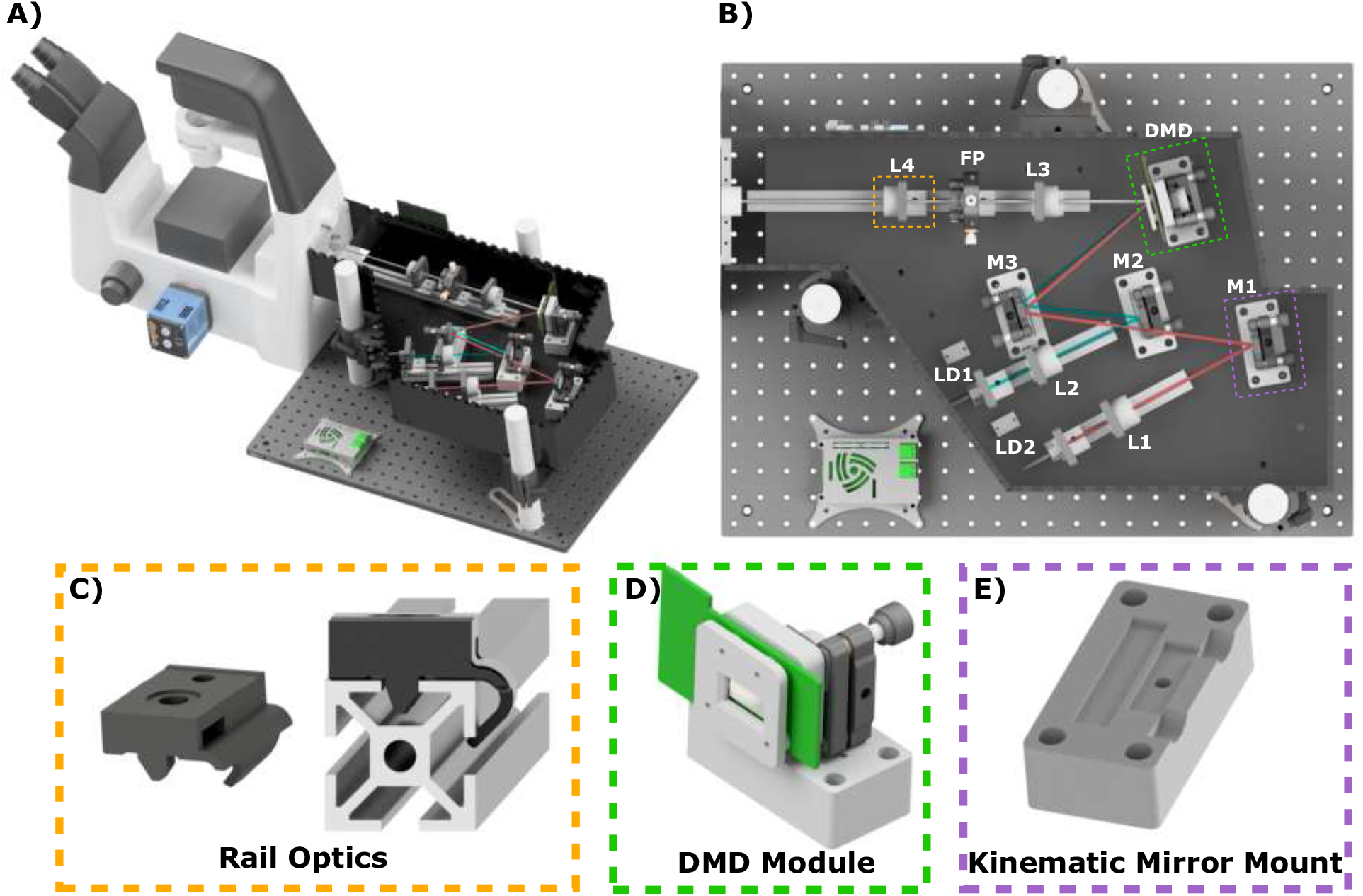
A) the complete openSIMMO setup attached to a Nikon Ti2-A Eclipse. B) The beam path of the two lasers is folded to minimize the footprint and meet the blaze condition. C) Aside from the laser-cut housing and a minimal number of external optics components, the mechanical design relies on the RailOptics system that enables translation of optics along the optical axis. D) the DMD module is positioned with a customized optomechanical mount and driven via the HDMI port of a Raspberry Pi. E) off-the-shelf optics are adjusted using 3D printed adapters mounted on a laser-cut base plate.

After the DMD, the illumination pattern is relayed by a 1:1 telescope consisting of two 75 mm focal length lenses (AC-254-75-A M, Thorlabs) and a Fourier mask is deployed to intercept the Fourier plane and obstruct the main diffraction order, leaving only the two subdiffraction orders for 2D SIM illumination. The mask was fabricated by drilling holes into black aluminium foil (Thorlabs BKF12), which is further described in the online documentation. A pizza polarizer, as proposed in [22], was not included because the increased contrast advantage was not outweighed by the increased cost and complexity of construction (i.e. due to the compact telescope, the small diameter of the Fourier plane makes it difficult to arrange individual quarter-wave plates). A tube lens (Thorlabs AC-254-200 A M) then focuses the sub-order beams on to the pupil plane of the microscope objective (60x/1.4 NA, Nikon, Japan), which is part of an inverted microscope stand (Ti2-A Nikon, Japan).

The patterned light excites fluorescence from the sample, which is recaptured by the objective lens, passes through a dichroic mirror (N-SIM 405/488/561/640, Nikon, Japan), and is focused onto the camera sensor (PCO Edge 4.2, Kehlheim, Germany) with a pixel pitch of 6.5 µm by a tube lens inside the microscope body. The effective pixel size is 108 nm, implying slight undersampling with regard to the Nyquist limit for the 488 nm excitation, and correct sampling for the 635 nm excitation, which would be 187 nm and 239 nm respectively. To fit the Nyquist criterion for the 488 nm excitation an in-built 1.5å intermediate magnification lens (IML) from the Nikon Ti2-A was employed during the experiment.

Alternatively, for some experiments, we used an industry-grade USB3 camera (Daheng, MER2-230-168U3M, China) to show the ability of cost-effective CMOS camera applied into scientific imaging. The Daheng camera uses an IMX174 sensor which has a pixel pitch of 5.8 µm and an effective pixel size of 97 nm. It matches the Nyquist sampeling criterion for 488 nm excitation with 60x microscope objective. In order to demonstrate the advantages of SIM in optical sections with a large field of view, we also used a lower magnification objective (20x/0.75 NA, Nikon, Japan) in addition to the super-resolution approach.

Our mechanical design integrates commercial parts, craft materials, and 3D parts to create a stable, adjustable, and affordable experimental platform. Specifically, we incorporate two principal types of mounting hardware. First, aluminium profiles — referred to as “RailOptics” — afford critical adjustability for components along the optical path, such as the laser collimation lenses and the telescope. These are attached to a 3D-printed part (see Fig. 1b-e) that clips onto the aluminium profile, allowing for precision movement along the optical axis and facilitating quick changes and adjustments. Second, specialized 3D-printed bases house kinematic mirror mounts (KM100, Thorlabs, US), provide fine angular adjustments to ensure the mirrors and the DMD chip are precisely aligned with the optical axis.

### C. Automation of the Image Acquisition and Reconstruction

OpenSIMMO is designed to achieve high-resolution SIM reconstructions in real-time during *in vitro* experiments. The hardware control, image acquisition, and data reconstruction must be orchestrated so that the high-resolution data can be displayed while live cell experiments are conducted. The open-source control platform ImSwitch controls the various hardware components and synchronizes the individual modules. A customized plugin extends this napari-centric [39], Python-based software with the option of entering SIM parameters and starting and controlling various experiments.

Here, openSIMMO benefits from a variety of different open-source algorithms for reconstructing the raw SIM images into a super-resolution result. napari-SIM [40, 41] supports reconstructing raw data stacks in real-time after prior calibration of the illumination pattern parameters. The result is displayed directly in the napari viewer. The calculation can be accelerated if a graphics card is available. The same applies to the mcSIM library [42], which is a Python package that can also be integrated with napari. Alternatively, the stacks can be imported into imageJ and reconstructed with fairSIM [16].

One difficulty is the lack of trigger input/output on the DMD evaluation board. An effective hardware-controlled and time-synchronous display of the previously calculated SIM patterns for the different lenses and wavelengths on the DMD was therefore not possible. For this reason, an additional control device, the Raspberry Pi 3b (Raspberry pi foundation, UK), was integrated into the workflow (Fig. 3). A REST-API-based display server streams the patterns stored on the device via HDMI to the DMD evaluation module and can simultaneously output trigger pulses for the camera. ImSwitch starts the continuous image acquisition via the “fastAPI” interface and sends the collected image stack containing images for all illumination patterns, wavelengths and focus positions to the reconstruction algorithms.

Compared to hardware trigger-based displays, the maximum possible recording speed is limited by the Raspberry Pi’s refresh rate of 60 Hz. However, the limiting factor in the experiments shown here was the required minimum exposure time of ≥ 50 ms set by the relatively weak lasers.

To control additional hardware components, such as the focusing motor and incubation unit, the UC2-REST system was used, which is fully accessible via a USB serial connection through the ImSwitch GUI [43]. The ESP32-based system enables TTL-based control of the laser intensity, as well as the realization of a heating chamber to perform live cell experiments. A heating plate from a 3D printer (Prusa Mini, Prag, Czech Republic) was mounted in conjunction with a temperature sensor in a 3D printed case on the sample stage. A PID controller running on the microcontroller regulates the temperature to a constant value of 37 ^°^C. To produce image stacks in focus, a NEMA17 motor was attached to the manual focus drive of the microscope by means of a belt drive.

Our system can be easily adapted to a wide variety of different harware due to the flexibility of ImSwitch. For example, ImSwitch offers a large variety of different camera drivers hardware adapters. In this work, work we have used this flexibility to profile openSIMMO using both scientific and industrial CMOS cameras.

## RESULTS

A primary goal of openSIMMO is to make super-resolution microscopy using SIM available to a large user base. The associated need to develop an affordable, compact, easy-to-build system composed of widely available parts has limited the space of possible designs and led to a slight loss in maximum resolution, since we, for example, omitted the use of the pizza polarizer, which is required to achieve maximum modulation contrast especially for high-NA objective lenses. We characterize the system as follows, both in terms of biological and synthetic calibration samples, regarding the obtainable lateral and axial resolution, as well as the ability to use multiple colors sequentially and to record time-lapse series.

Here we worked with pattern periods of 310 nm for the 488 nm excitation and 425 nm for the 635 nm excitation (section S4). These patterns support increasing the maximum detectable spatial frequency by a factor of ≈1.55, ignoring the effect of the Stoke’s shift.

First, the maximum resolution of the setup was measured with a SIM ArgoLight slide (Argolight SIM v1, SLG-008, Argolight, France) using the 488 nm excitation. The finest line pair that can be resolved with the deconvolved widefield image is 210 nm. After reconstructing the data, the line pairs with a distance of 120 nm can be clearly resolved. (see Fig. S1 A,C). This corresponds to a 1.75-fold improvement compared to the Abbe diffraction limit, whereby a 1.55-fold improvement would have to be expected. The difference could be due to the undersampling with 488 nm laser excitation without using the in-built 1.5å intermediate magnification lens. For getting higher signal of the raw data, the magnification lens was removed to increase the photon number on each camera pixel.

To characterize system multicolor performance with biological samples, we imaged a fixed huFIB cells sample with multiple fluorescence markers (GATTA-Cells 4C, Gattaquant, Germany) and the results are shown in Fig. 4 A,E. Here, the mitochondria are labelled with Alexa Fluor 488 and imaged with the 488 nm excitation, while the actin is labelled with Alexa Fluor 647 and imaged with the 635 nm excitation. In addition to the clear increase in resolution, the improved optical sectioning should be emphasized. The fluorescent background is suppressed using Gaussian zero-order suppression (default parameters for Napari-SIM and FairSIM were α = 0.5, β = 0.98 and a = 0.99, FWHM = 1.2), leading to an improved imaging of the cell structure. The cell image resolution in the 488 nm and 635 nm channel are validated with the Fourier-ring correlation (FRC) method [44] which stimates a resolution of 130 nm vs. 220 nm and 174 nm vs. 260 nm for the pseudo widefield image. A 3D stack can be done with moving the employed NEMA17 motor which is attached on the fine focus knob. The stack was captured with 260 nm spacing over total range of 10.4 µm as show in Fig. 3B.

Our open-source SIM module is the centerpiece of a low-cost, open-source live cell imaging platform. The open-source nature of ImSwitch enables us to implement additional features such as autofocus which are vital for live-cell imaging and time-lapse recordings. We demonstrate these functionalities on the recording of living cells, which is realized using a self-built incubator module. A PID controller implemented in UC2-ESP firmware [45] ensures a constant temperature of 37 ^°^C, enabling affordable *in vitro* imaging of living cells.

We demonstrate our instrument’s live cell imaging capabilities by performing a timelapse series of HeLa cells cultured with MitoTracker Green dye. Fig. 3C shows two of the total 30 time points (*t*_period_ = 2 min, *t*_exposure_ = 50 ms), where moderate bleaching and a focus drift of the objective of about 20 µm*/*h was observed. The fine focus knob was turned mechanically by the motor before each image series acquisition to balance the focus drift. For the future it could be compensated using an autofocussing algorithm implemented in ImSwitch. Compared to the widefield, a much better optical sectioning is observed in the SIM reconstruction (Fig. 4). The resolution of the time-lapse widefield and SIM data are estimated to be 226 nm and 158 nm by using FRC, which gives a resolution enhancement of 1.43å.

For the time-lapse and 3D stack experiments presented here, we used the 60å objective in combination with the Edge 4.2 sCMOS camera without the intermediate magnification lens and the data is ≈10 % undersampled at the ≈520 nm emission wavelength corresponding to the 488 nm excitation. Since the OTF rapidly falls off at high spatial frequencies, we did not observe aliasing effects in the raw or reconstructed data.

## DISCUSSION

In this paper, we introduce an illumination add-on specifically designed for structured illumination microscopy attached to a standard commercial fluorescence microscope body. Our configuration is enclosed within a laser-cut Plexiglas box, where components are mounted onto the baseplate similar to commercial illumination add-ons. The add-on is fully opensource, including a Python-based framework that has access to a large variety of different cameras and hardware control elements using napari and ImSwitch. We demonstrate that our setup, which in its minimal form incorporates an industry-grade camera, off-the-shelf motors and microcontrollers, and a DMD-based SIM module, enables a powerful SIM upgrade to existing microscopes for approximately €3000, democratizing access to superresolution imaging capabilities. Additionally, our adaptable control software supports various hardware configurations, potentially revitalizing outdated microscopes into state-of-theart super-resolution devices for live cell experiments, thus giving scrap-heap equipment a second life. Our implementation simplifies the integration of super-resolution imaging into fully automated workflows, where e.g. automated sample preparation can be part of a larger pipeline. A remote-controllable REST-API integrates triggering of super-resolution imaging and reconstruction in a unified way. Every part of this project has been open-sourced, allowing for collaborative contributions. A step-by-step demonstration of preparing, building, aligning, and starting the extension setup enables widespread adoption.

By minimizing the use of commercial opto-mechanical parts and by using 3D printed parts instead, we have increased the reproducibility. Through a series of rigorous experiments, we consistently obtained reliable results. Our finalized setup enhances resolution, surpassing that of a traditional widefield microscope by factor of 1.55. This extension can be effortlessly affixed to microscope bodies with an epi-fluorescence port, enhancing its versatility. The accompanying software is accessible to users of all levels of expertise.

While openSIMMO achieves excellent balance between low-cost and high-quality superresolution imaging, some of the hardware compromises limit some aspects of the setup’s performance. In order to get an acceptable result in the current configuration with the selected lasers of limited power, the exposure time for each frame needs to be longer than 50 ms, hence limiting its use for rapidly moving objects in live-cell imaging. The speed limit of the DMD has thus not yet been reached. An upgrade to a high power laser holds the potential to further enhance the setup’s operation speed, further pushing the boundaries of its performance. Improving polarization control has the potential to improve imaging contrast (further quantified in Sec. S5) and thus the maximum achievable resolution improvement. For example, the implementation of circular polarization, e.g. with quarter-wave plates (Thorlabs, WPQSM05-X approx. 300 euros), is a cost-effective solution for improving contrast if the budget permits.

Alternatively, the existing openSIMMO design could be made more affordable by adapting widely used projectors from DLP 3D printers could provide a powerful but even cheaper alternative for pattern generation and offer the possibility of triggering the camera, as recent work has shown [46].

We anticipate that openSIMMO will democratize super-resolution fluorescence imaging and significantly expand the availability of high-contrast coherent SIM instruments in the life sciences. We believe broader dissemination of this technology will have major impacts on fluorescence live cell imaging and improve the quality and reproducibility of imaging experiments generally.

## Supporting information

Explosion View of CAD Assembly of SIMMO

## ACKNOWLEDGMENTS

We acknowledge funding from the Nexus GIF grant no. G-1566-143.13/2023. We thank the free state of Thuringia for supporting this project. We acknowledge Frank Garwe for his assistance in the grant application and Gerhard Holst for constructive feedback on the camera. We also thank Nikon for allowing us to use an instrument to which our setup was attached to.

## DATA, MATERIALS, AND SOFTWARE AVAILABILITY

All data associated with this manuscript are available in our GitHub repository https://opensimmo.github.io/

## AUTHOR CONTRIBUTIONS (IN ALPHABETICAL ORDER)

Conceptualization: B.D., H.W., P.B. Data curation: H.W., P.B.

Funding acquisition: B.D. Investigation: B.D.

Methodology: B.D., H.W., D.P.S., J.U., P.B., R.H.

Project administration: B.D.

Resources: B.D., D.P.S., R.H.

Software: B.D., H.W., P.B. Supervision: B.D. Validation: B.D., H.W., P.B.

Visualization: B.D., H.W., P.B.

Writing – original draft: B.D., H.W., P.B., R.H. with input from all authors.

Writing – review & editing: All authors

## SUPPLEMENTAL INFORMATION

## Appendix S1 Sample Preparation

HeLa cells were cultured in Dulbecco’s Modified Eagle Medium (DMEM) supplemented with 10 % Fetal bovine serum (FBS) and 1 % penicillin/streptomycin solution (P/S) and maintained in a 37^°^C/5 % CO2 incubator. Cells were harvested around 90% confluency with TrypLE express 1x (12604013, ThermoFisher Scientific, Germany) and transferred into ibidi 4 well slide (µ-Slide 4 well, ibidi GmbH, Germany), the sample slide is cultured again for one day to make sure cells adhere to the glass bottom.

For live cell imaging, cells were stained with MitoTracker Green (Invitrogen MitoTracker Green, ThermoFisher Scientific, Germany) and imaged using 488 nm laser. To prepare the staining solution, 1 µL MitoTracker Green stock solution was diluted into 1 mL cell culture medium in order to produce staining solution with concentration of 1 µM. When the cells achieve 70%-90% confluency, remove the old culture medium and wash the sample with PBS three times, then add the staining medium to the slide and culture the sample in the incubator for 1 h before imaging.

When the HeLa cells in ibidi slide multiply up to 70%-90% confluency, the sample was fixed with 4 % Paraformaldehyde solution (PFA) for 15 min at room temperature. After fixation, the sample was washed with PBS three times in order to remove the PFA solution totally. 0.1 % Triton X-100 diluted in PBS was used to permeabilize the cell membrane for 2 min. 2 µL Alexa Flour 488 Phalloidin stock solution was diluted with 400 µL PBS as staining solution for each well. The sample was stained with staining solution for 50 min at room temperature and later stored in PBS solution.

## Appendix S2 Lineplot of Argolight calibration sample

**FIG. S1.**
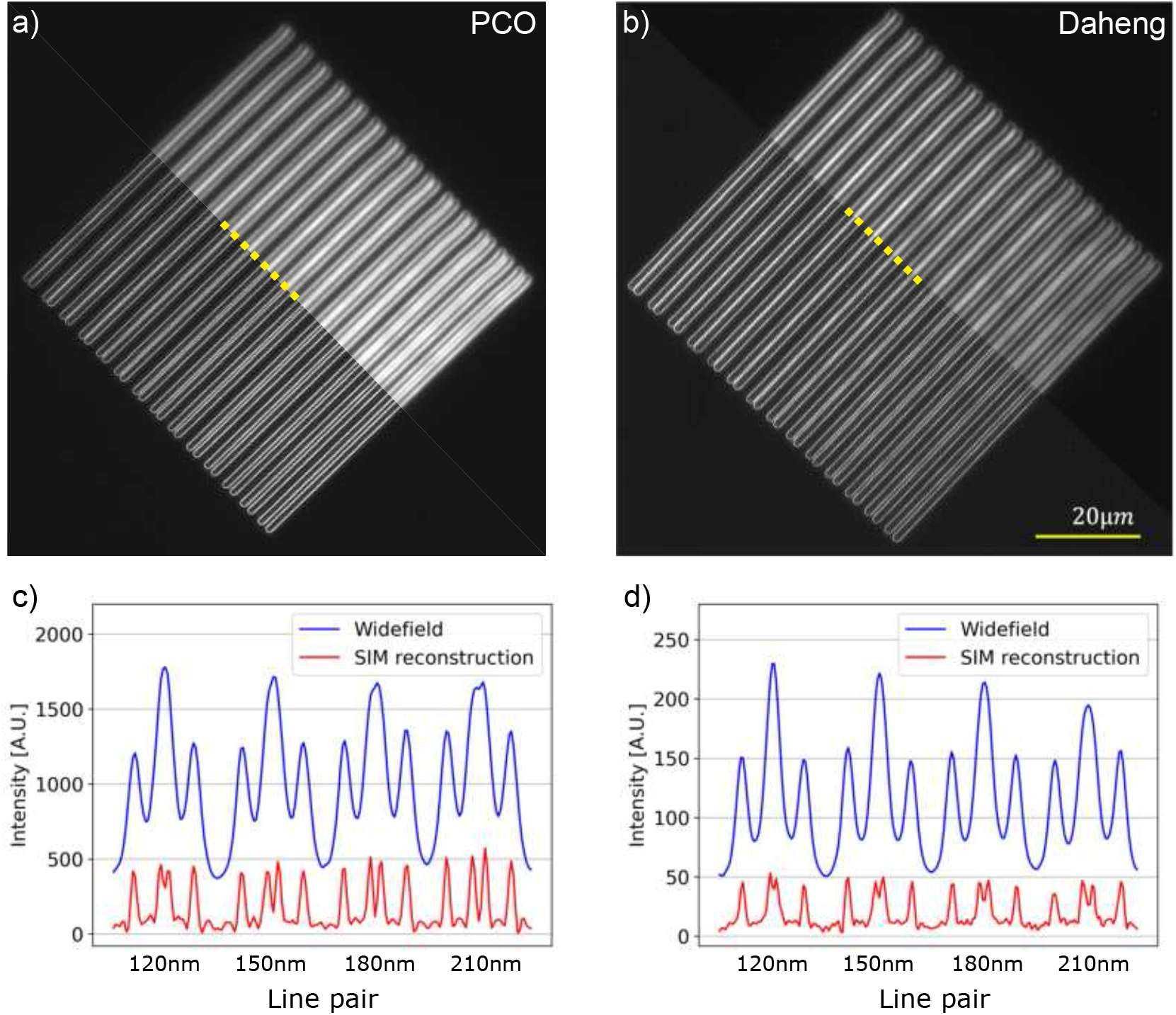
Resolution characterization of the system with different cameras including the intermediate magnification lens. The data show the line pair on the Argolight SIM calibration slide excited with a 488 nm laser. Wide-field and SIM results with a) the PCO Edge 4.2 camera and b) the Daheng MER2-230-168U3M camera. c) and d) show the line plot of the line pair between 120 nm and 210 nm. The line plot data is averaged over 5 pixels.

## Appendix S3 Imaging with industrial camera

For a cost-effective solution, an industrial grade CMOS camera (MER2-230-168U3M, Daheng,China) was tested on the setup. The camera has a CMOS chip IMX174 with pixel pitch of 5.86 µm

**FIG. S2.**
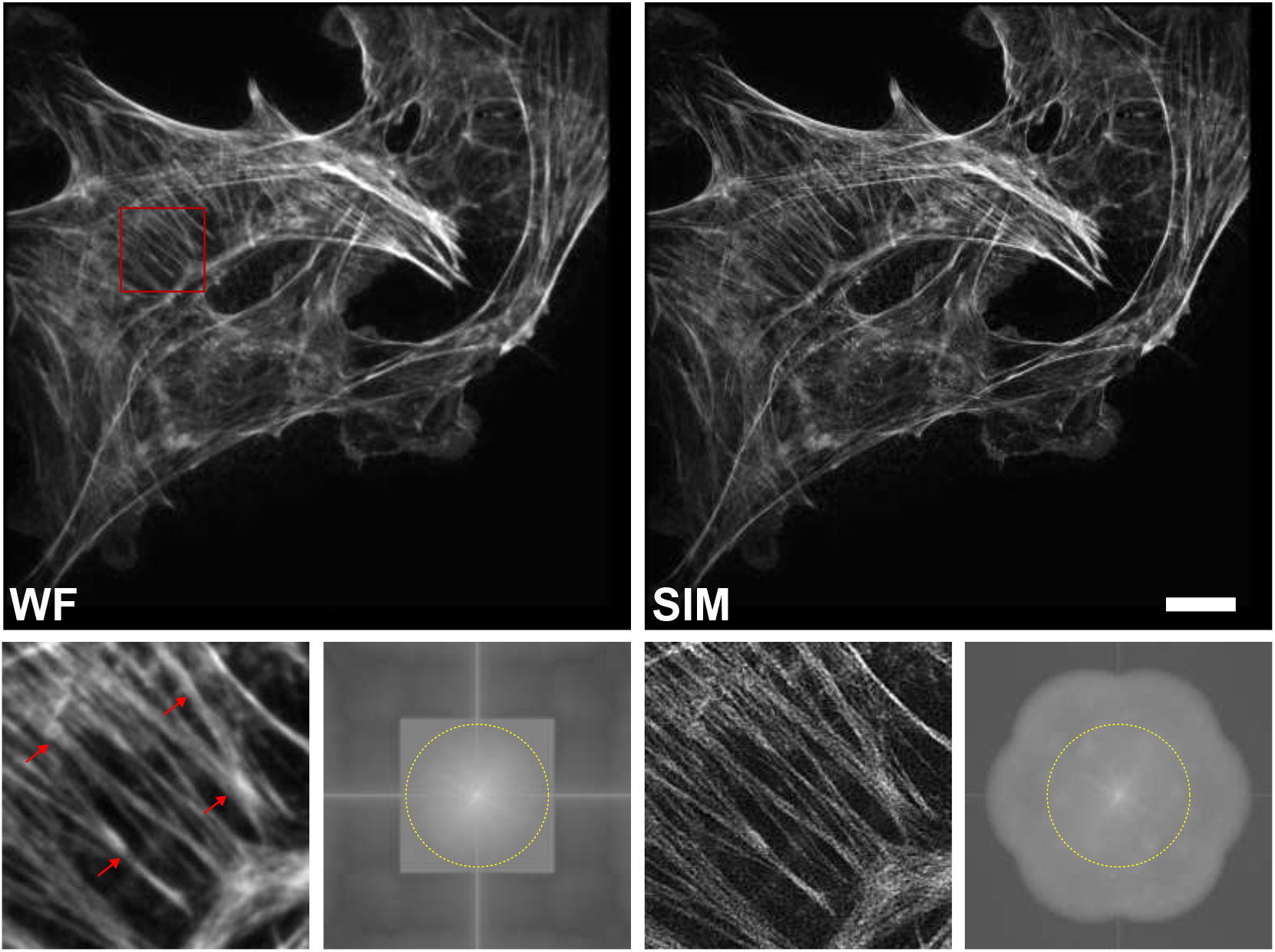
BPAE cells from a FluoCells Nr. 1 slide were imaged using the Daheng camera. We compare the widefield image (upper left) and SIM reconstruction (upper right), SIM reconstruction gives better resolution compared to the widefield. To illustrate the enhanced SIM resolution we display a region of interest (lower left and third from left). To illustrate the increased frequency support in Fourier space, we also display the Fourier transforms of the widefield image (lower, second from left) and SIM reconstruction (lower, right). Scale bar is 10 µm.

**FIG. S3.**
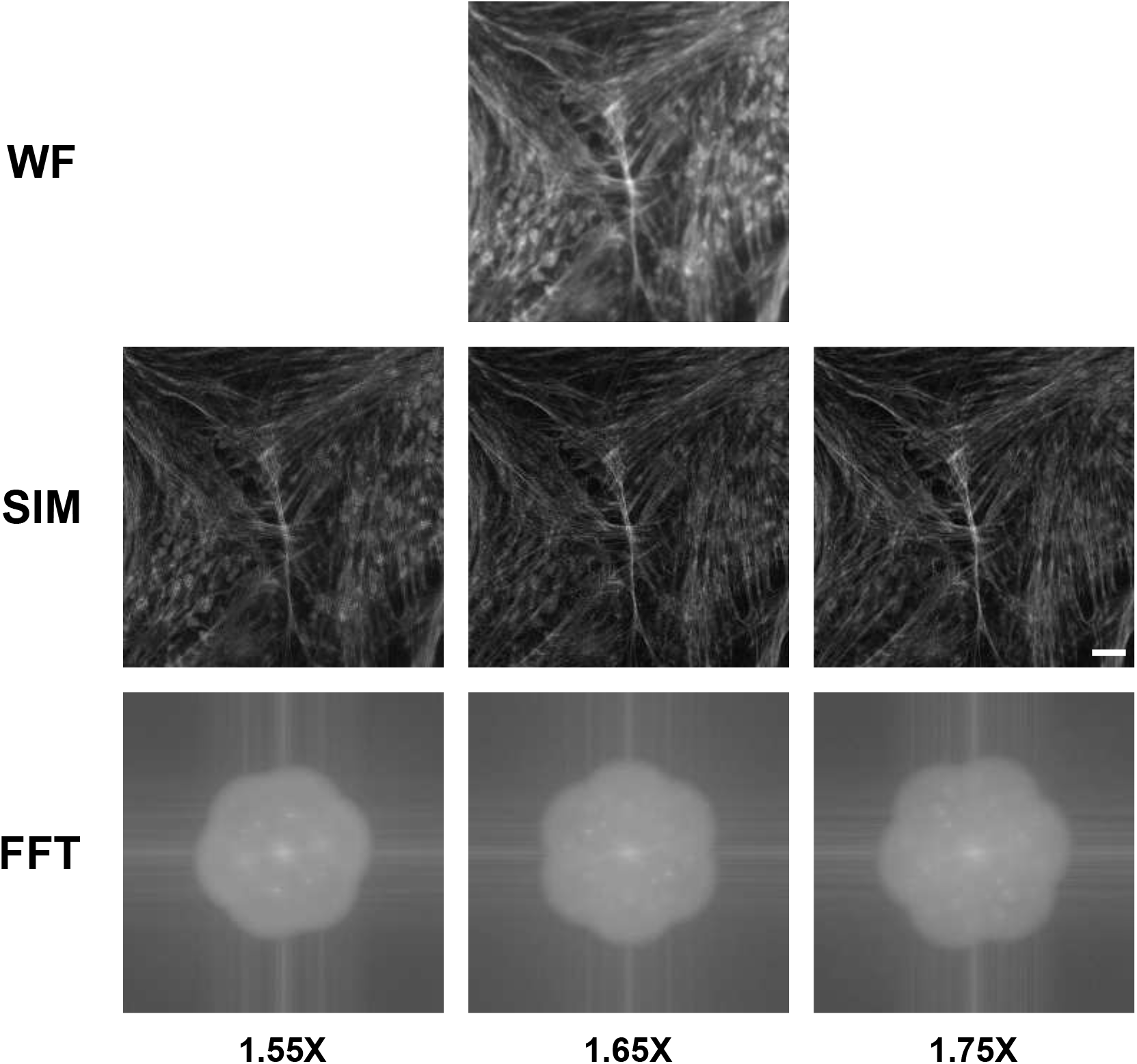
Different patterns were used in the experiment to see the difference between varied grating constant for the resolution enhancement. The sample shown here is stained BPAE cells on FluoCells Nr. 1 and excited with 488nm laser. The results show with finer grating applied to the sample, the resolution of the image is slightly increased, which is also proved on the Fourier transformation of the data with more higher frequency information. Data were captured with PCO Edge 4.2 camera. Scale bar is 5 µm.

**TABLE S1.**
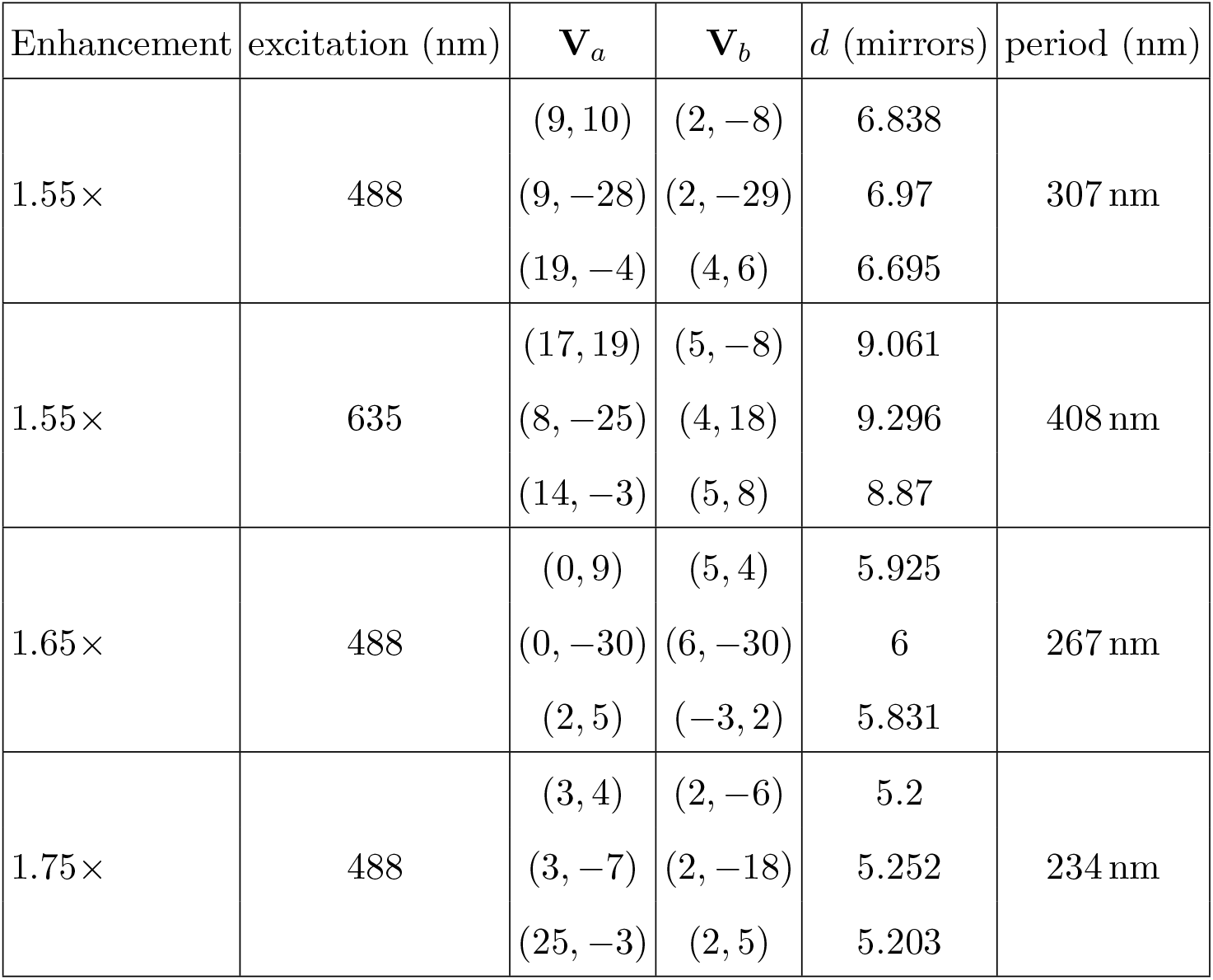
DMD SIM pattern parameters for the various patterns considered in this work.

## Appendix S4 SIM Pattern Generations

The DMD patterns used to project the SIM illumination were generated using the fastSIM grating search algorithm [14] implemented in fairSIM [16]. For different experiments, we used SIM patterns designed to provide resolution enhancements of 1.55, 1.65, or 1.75 times. The DMD pattern parameters are provided in table S1. In order to capture the pattern information with smaller grating constant an in-built 1.5å intermediate magnification lens (IML) in the Nikon Ti2-A Eclipse was employed in the imaging path. In this work, we used the 1.55 å patterns for the dual color data presented in the main manuscript. We used 1.65å and 1.75å patterns for 488 nm excitation in Fig. S3.

## Appendix S5 Effect of Polarization on Pattern Contrast

In coherent SIM, the control of polarization directions critically influences the intensity contrast of the illumination patterns. Perfect interference occurs when the light is polarized orthogonal to the plane formed by the beam propagation direction and the optical axis, while for light polarized parallel to this plane the contrast *m* is reduced to cos(2θ), where θ is the angle between the incident SIM beams and the optical axis. Note that the contrast *m* is negative for θ *>* π*/*4, just indicating a flip of the pattern phase. Averaging over various input directions, such as with circularly or unpolarized light, the modulation contrast becomes 0.5 (1 + cos(2θ)) [27]. Although linearly polarized light can provide excellent interference at one SIM angle, it may perform poorly at others. If the polarization cannot be controlled separatley for different SIM angles, circularly polarized light is more advantageous, as ‘half’ of it interferes perfectly.

In reality, the light exiting the single-mode fiber coupled lasers is neither linear nor circularly polarized leading to a maximum achievable pattern contrast of 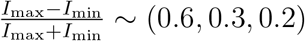 for the three different rotational angles for SIM pattern period 310 nm, NA = 1.4, and the 488 nm excitation. These values were calibrated with a sparse bead sample (TetraSpeck microspheres, 0.1 µm, Thermo Fisher, MA, USA). This contrast, although reduced, still surpasses that achievable with incoherent SIM, thereby enhancing resolution reconstruction.

## Appendix S6 DMD model

### A. Blaze condition optimization

Here we discuss the solution to the DMD diffraction optimization problem described in the main text. We have 8 unknowns, including the Lagrange multipliers, and 8 equations. However, since the optimization problem only depends on the vector difference 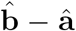, we will find one of these equations is redundant.

To evaluate the derivatives of the cost function, we use the basis change expressions,

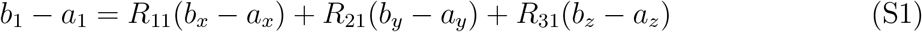

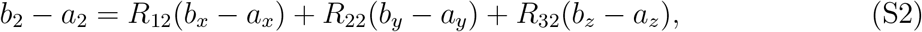

Then equation 8 becomes

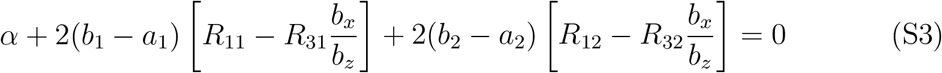

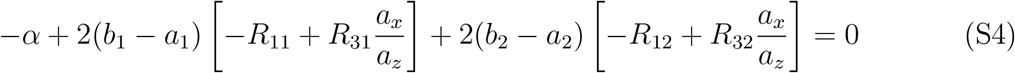

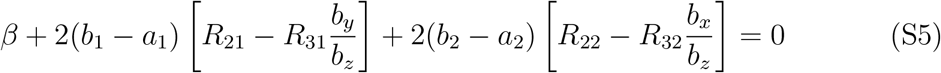

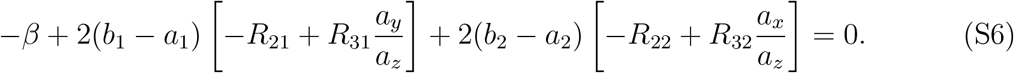

Adding eqs. S3 and S4 and separately S5 and S6 gives

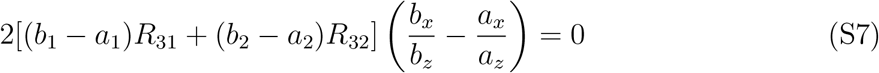

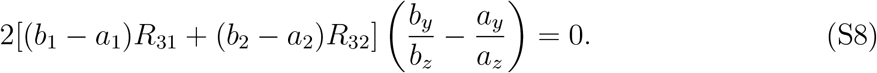

This leaves two possibilities (i) both terms in round brackets are zero, implying 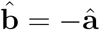 and the solution is unique or (ii) the term in square brackets is zero, and there is a one-parameter family of solutions.

In case (ii) we have

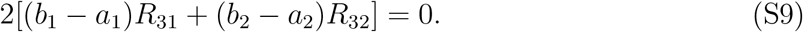

Substituting eqs. S1 and S2 into eq. S9, we find,

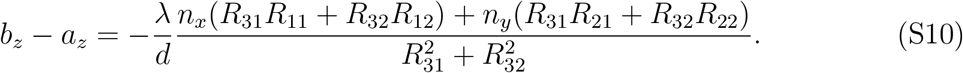

Combining this result with the diffraction condition fixes the value of 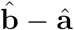 In principle, we also know the values of the Lagrange multipliers by reinserting eq. S10 into eqs. S3–S6. We have three parameters left to fix, which are the components of either â or 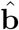. However, since the cost function only depends on the difference vector, this will not affect its value. Therefore, we have full freedom to choose one unit vector component, say *a*_*x*_, and we will still have a solution to our equations. That is, our system of equations is actually underdetermined because two of the Lagrange multiplier equations were identical.

Therefore, it remains only to express *a*_*y*_ in terms of *a*_*x*_ and 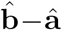 -â. One convenient approach is to notice that combining eqs. 3, 4, and S10 also fixes 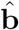 · â in terms of the DMD geometric parameters and the diffraction order

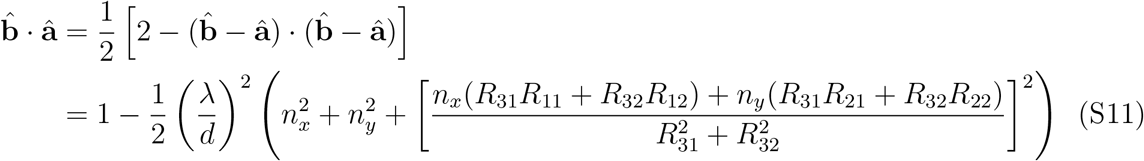

We now derive a quadratic equation for *a*_*y*_ in terms of *a*_*x*_, where all other vector parameters are expressed in terms of *a*_*x*_, the chosen diffraction order, and the DMD geometric parameters. To do this, we rewrite the dot product in terms of components, eliminate the *z*-components using eq. 1 and eq. 2, and square both sides

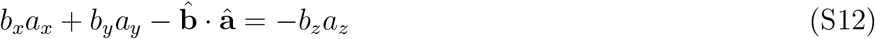

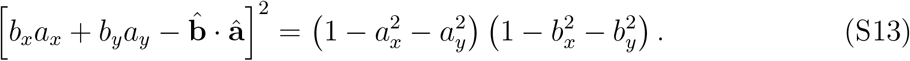

Finally, we rewrite eq. S13 in the form

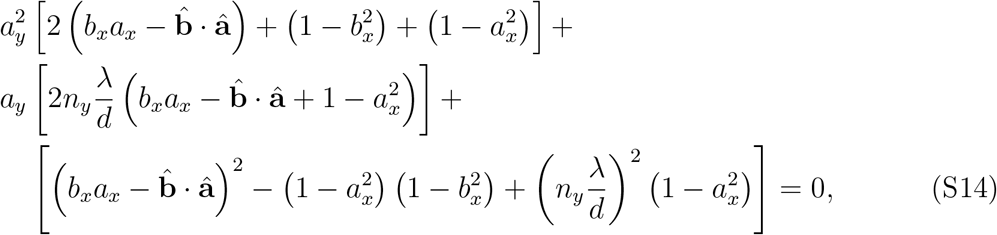

where we have explicitly eliminated *b*_*y*_ by inserting eq. 4 and implicitly eliminated *b*_*x*_ and 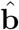 · â using eqs. 3 and S11 respectively. This quadratic equation can now be solved for *a*_*y*_ as a function of *a*_*x*_. Since eq. S14 was obtained by squaring eq. S12, we must check that the resulting solutions also satisfy eq. S12. Those that do are the final solutions to our optimization problem and recover the exact solutions to the combined diffraction and blaze conditions, if they exist.

**FIG. S4.**
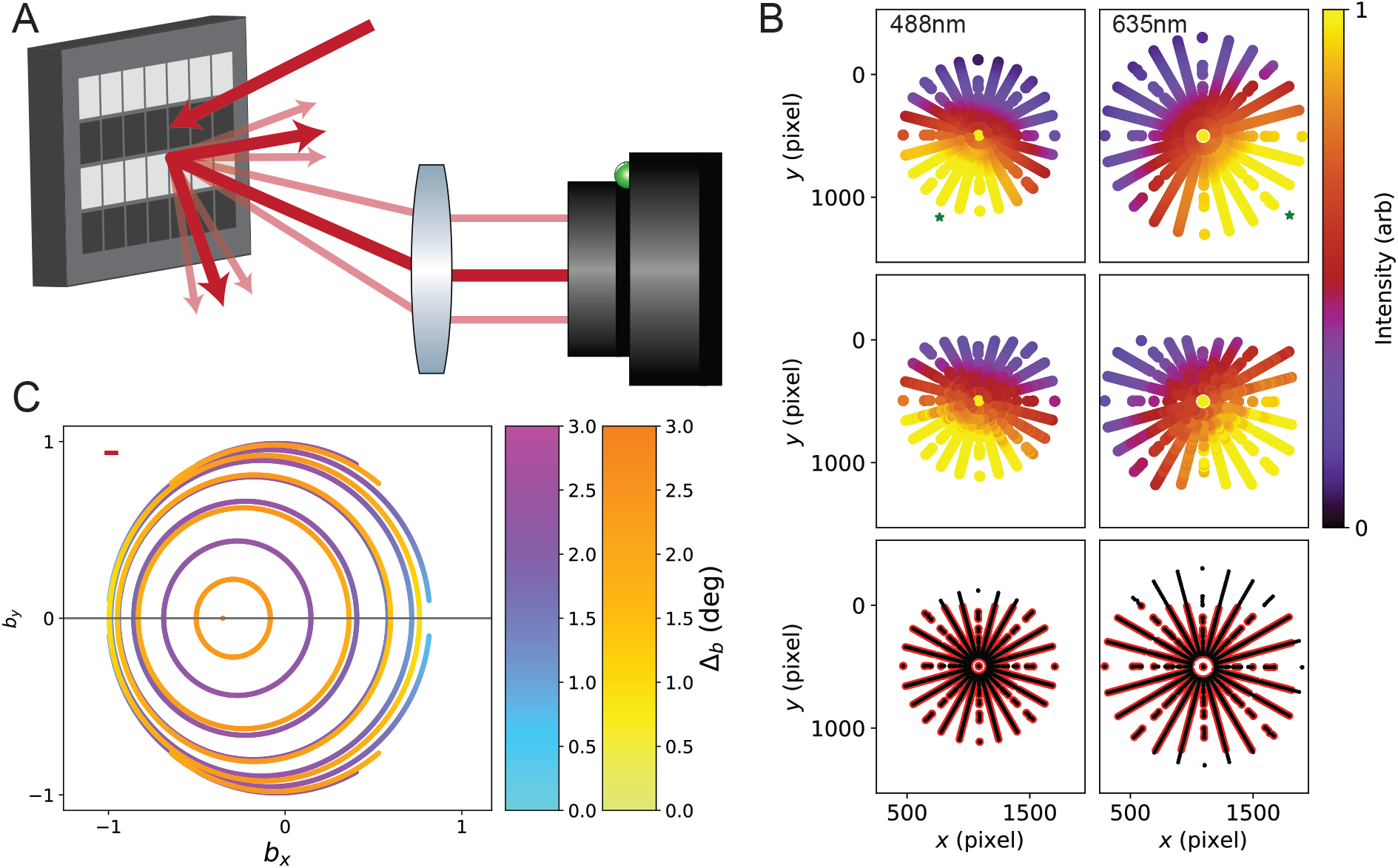
A). Coherent light incident on the DMD pattern is diffracted into many orders, and the intensity and position of some of these orders are measured on a camera placed after a lens in a 2*f* -configuration. B). We compare the predictions of our DMD diffraction model (top row) to the experimental measurements (second row) after applying an optimization procedure to determine the most likely beam geometry and DMD parameters. The predicted centers of the blaze envelope are shown (green stars) C). We recompute the optimally blazed DMD solutions using the recovered parameters, and find that due to deviations between the nominal and actual mirror rotation, the achievable blaze angle violation is a factor of ≈3 worse than expected. We consider diffraction orders (−*n*, 0) for *n* = 1, …, 6. For 488 nm, the innermost closed curve corresponds to the (−6, 0) order. For 635 nm, the (−6, 0) order solution is a single point, and the first closed curve corresponds to (−5, 0).

### B. Pixel Rotation Matrices

We can express any rotation matrix as a rotation through angle γ about axis 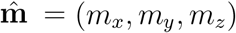 In this case, the rotation matrix is given by

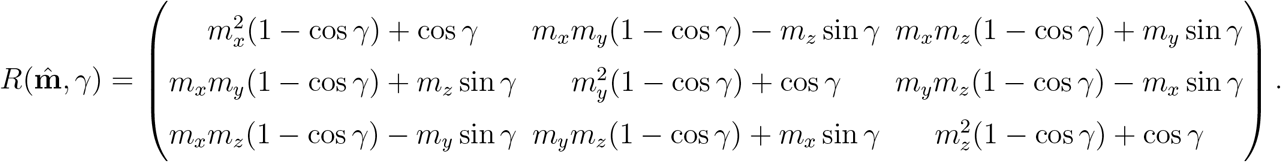

### C. Corner Illumination Pixels

For CIP’s, the rotation matrix parameters are

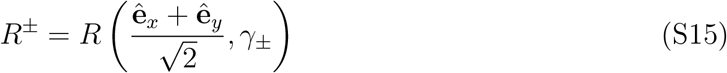

with γ_±_ = ±12°. Since *R* = −*R*, we only have exact solutions for *n*_*x*_ = −*n*_*y*_ = *n*, and in this case eq. 6 implies

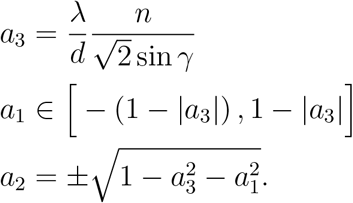

### D. Tilt and roll pixels

The TRP’s use a mechanism that tilts the mirror first by 12° degrees along one diagonal, then rolls it ±12° along the other diagonal to reach the + and states, respectively. The + position is tilted approximately 17° along the *y*-direction, while the position is tilted approximately -17 degrees along the *x*-direction [47]. Rewriting each composite rotation along a single axis,

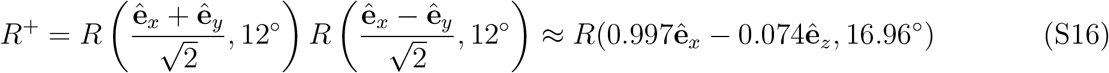

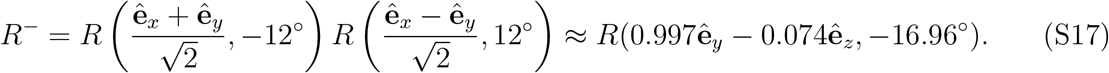

**TABLE S2.** DMD parameters inferred from validation experiments compared with expected values.

| Parameter | Expected Value | Fit Value | Comments |
| --- | --- | --- | --- |
| Blue diffraction order | (-6, 0) |  |  |
| Red diffraction order | (-5, 0) |  |  |
| DMD pitch ($\mu\text{m}$) | 5.4 | 5.4 | Constrained to $\pm 0.001 \mu\text{m}$ of guess |
| $\lambda_1$ (nm) | 488(1) | 488.0(2) | Constrained to [487,489] |
| $\theta_{x,1}$ ($^\circ$) | 37 | 38.2(3) | $\hat{\mathbf{a}}_1 = (0.618, -0.003, -0.786)$<br>$\hat{\mathbf{b}}_1 = (0.076, -0.003, 0.997)$<br>$\Delta_b = 2.5^\circ$ |
| $\theta_{y,1}$ ($^\circ$) | 0 | -0.2(3) | |
| $\lambda_2$ (nm) | 637(2) | 635.0(3) | Constrained to [635,639] |
| $\theta_{x,2}$ ($^\circ$) | 40.5 | 41.7(3) | $\hat{\mathbf{a}}_2 = (0.665, -0.003, -0.747)$ |
| $\theta_{y,2}$ ($^\circ$) | 0 | -0.2(3) | $\hat{\mathbf{b}}_2 = (0.077, -0.003, 0.997)$<br>$\Delta_b = 3.2^\circ$ |
| $\theta_{x,p}$ ($^\circ$) | 3.4 | 6.7(8) | Lens optical axis orientation |
| $\theta_{y,p}$ ($^\circ$) | 1.6 | 1.5(7) | $\hat{\mathbf{p}} = (0.116, 0.026, 0.993)$ |
| $v_x$ (pix) | 1093 | 1773(220) | Camera affine transformation $x$-offset |
| $v_y$ (pix) | 501 | 998(160) | Camera affine transformation $y$-offset |
| $\theta_{\text{cam}}$ ($^\circ$) | 0 | -0.18(2) | Camera affine transformation rotation |
| $f_l/d_{xy}$ | 17 064 | 17 101(10) | $f_l = 100 \text{ mm}$, $d_{xy} = 5.86 \mu\text{m}$ |
| $f$ | 1 | 0.670(1) | Correction factor for DC peak strength |
| $w$ ($\mu\text{m}$) | 5.4 | 5.3 | Constrained to $w \geq 5.3 \mu\text{m}$ |
| $\theta_{\text{rot}}$ ($^\circ$) | 94.25 | 81(8) | $\hat{\mathbf{n}}_{\text{rot}} = (0.045, 0.987, 0.152)$ |
| $\phi_{\text{rot}}$ ($^\circ$) | 90 | 87(2) | |
| $\gamma$ ($^\circ$) | -16.955 | -17.7(2) | |
| $\alpha$ | 0.001 | | Relative strength of the position vs. intensity loss |

### E. DMD Model Validation

We validate our modelling approach by displaying different SIM-like patterns on the DMD and measuring the intensity and position of the resulting diffraction orders on a camera placed behind a lens in a 2f-configuration with the DMD [27] (Fig. S4A). For each pattern, we determine the position and intensity of the five most prominent diffraction orders by fitting each peak to a 2D Gaussian. We normalize each intensity by the DC peak’s intensity. We additionally normalize each peak intensity to the expected diffracted intensity based on the strength of each Fourier mode in the DMD pattern. Then, we compare the resulting normalized intensities and positions to the values predicted by our DMD diffraction model (Fig. S4B). We fit the model using a non-linear least-square approach to determine the beam input angles and wavelengths, the DMD mirror grid and mirror rotation parameters, and the imaging system alignment parameters. To obtain a robust parameter uncertainty estimation, we apply a bootstrapping procedure using 1000 samples [48]. We report the bestfit parameters using the full dataset together with the standard deviation of the parameters as determined using the bootstrap in table S2.

For the fit, we use different parameterizations of the parameters than elsewhere. For example, we describe the input directions in terms of angles *θ*_*x*_ and *θ*_*y*_, where

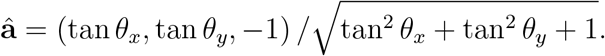

This is convenient, as these two angles are independent and have an unbounded domain, simplifying the optimization compared with working with unit vector components. Also, the parameterization ensures *a*_*z*_ *<* 0, so the represented vector is incident on the DMD. Alternatively, for the rotation axis, we adopt the parameterization

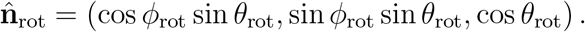

This again provides two unbounded independent variables but supports unit vectors with any sign for their z-component.

We find that the inferred DMD model parameters differ somewhat from the nominal values. The rotation axis of the - mirrors is oriented about 13° different than expected, and the mirror rotation angle is −17.7, slightly larger than expected. These parameters imply that the best achievable performance for 2-color operation with this DMD is somewhat worse than predicted for the nominal values (see fig. S4 C). In particular, the predicted blaze angle violations are on the order of approximately 3°, compared with approximately 0.5° as found in Fig. 2E.

**FIG. 2.**
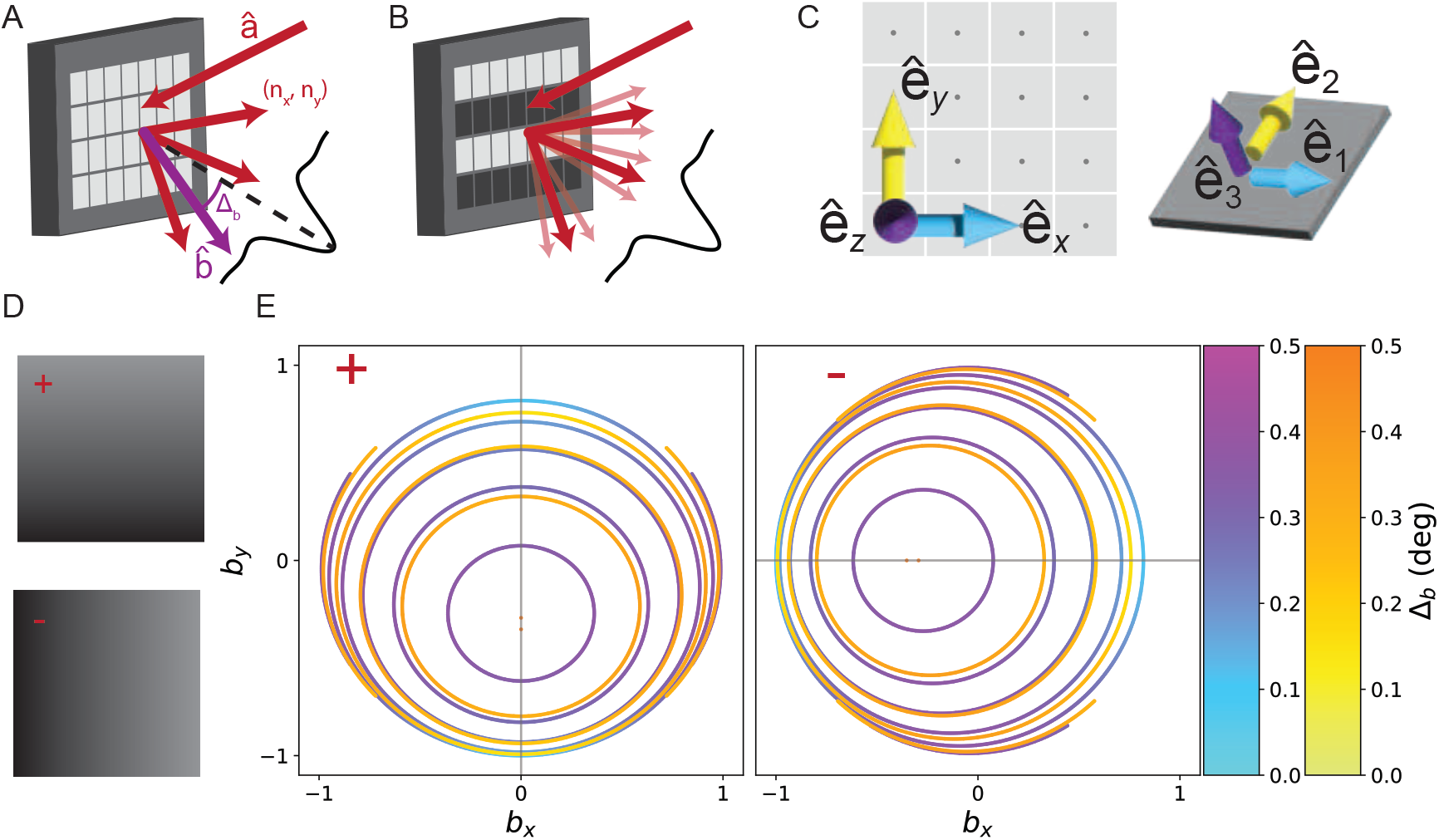
A). Coherent light incident on the DMD from direction â is diffracted into orders (*n*_*x*_, *n*_*y*_) whose strengths are modulated by the blaze envelope (black curve). We determine which output directions 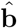 (purple) are allowed (red), and how much they deviate from the center of the blaze envelope, Δ_*b*_. B). When a pattern is displayed on the DMD, the additional structure produces subdiffraction (light red) centered around the primary diffraction orders (red). C). We define one basis aligned with the DMD backplane ê_*x*_, ê_*y*_, ê_*z*_ and a second basis which is adapted to the tilted mirror ê_1_, ê_2_, ê_3_. The two bases are related by a mirror-state dependent rotation matrix *R*^±^(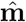, γ). D). TRP pixels support two states, “+” and “-” (left). The mirror rotation axis nearly coincides with ê_*x/y*_ for the ± states, respectively. The darker corner of the micromirror is the landed side. E). For each given diffraction order, many different input/output direction pairs lead to optimal solutions. We display these as curves in output direction space, parameterized by unit vector components *b*_*x/y*_. We determine the most nearly blazed diffraction solutions for 488 nm (blue to purple) and 635 nm (yellow to orange) incident light for several different diffraction orders for both the ± (left/right). For the + mirrors, we consider diffraction orders (0, *n*). For 488 nm, the innermost curve represents the (0, -6) order, and increasing area corresponds to lower order. For 635 nm, the first curve corresponds to (0, -4). The - mirrors show identical structure under a 90° rotation.

**FIG. 3.**
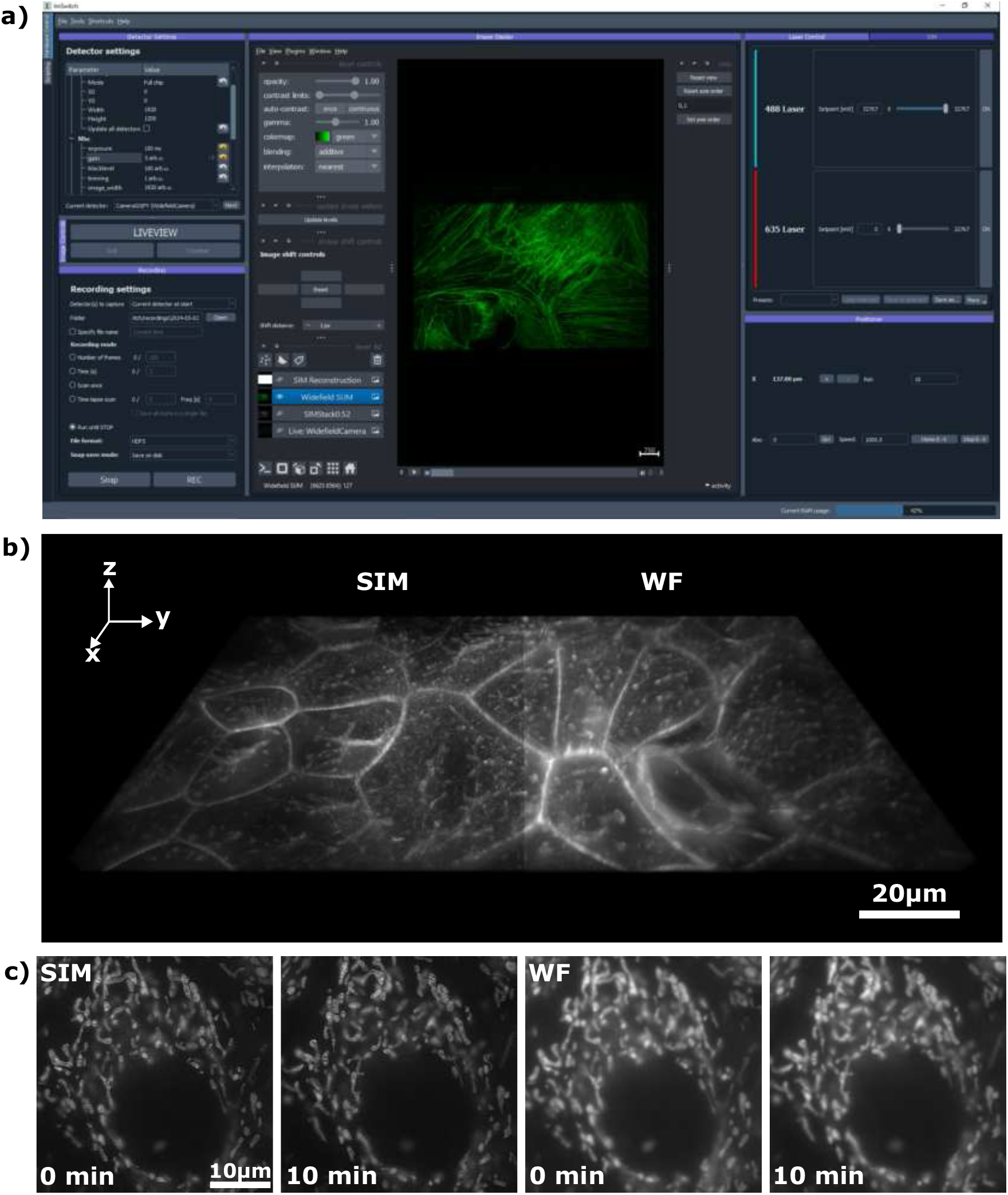
A) Integration of real-time SIM reconstruction into the open-source microscopy control, acquisition, and processing software, ImSwitch. The installable SIM plugin allows users to input additional experimental parameters, managing pattern display and camera frame acquisition across different manufacturers. The resulting images are displayed as layers in napari and can directly be stored to the disk. B) 3D stacks and C) time-lapse data, exemplified here with Mitotracker-labeled HeLa cells. The enhanced optical sectioning compared to widefield images is clearly evident.

**FIG. 4.**
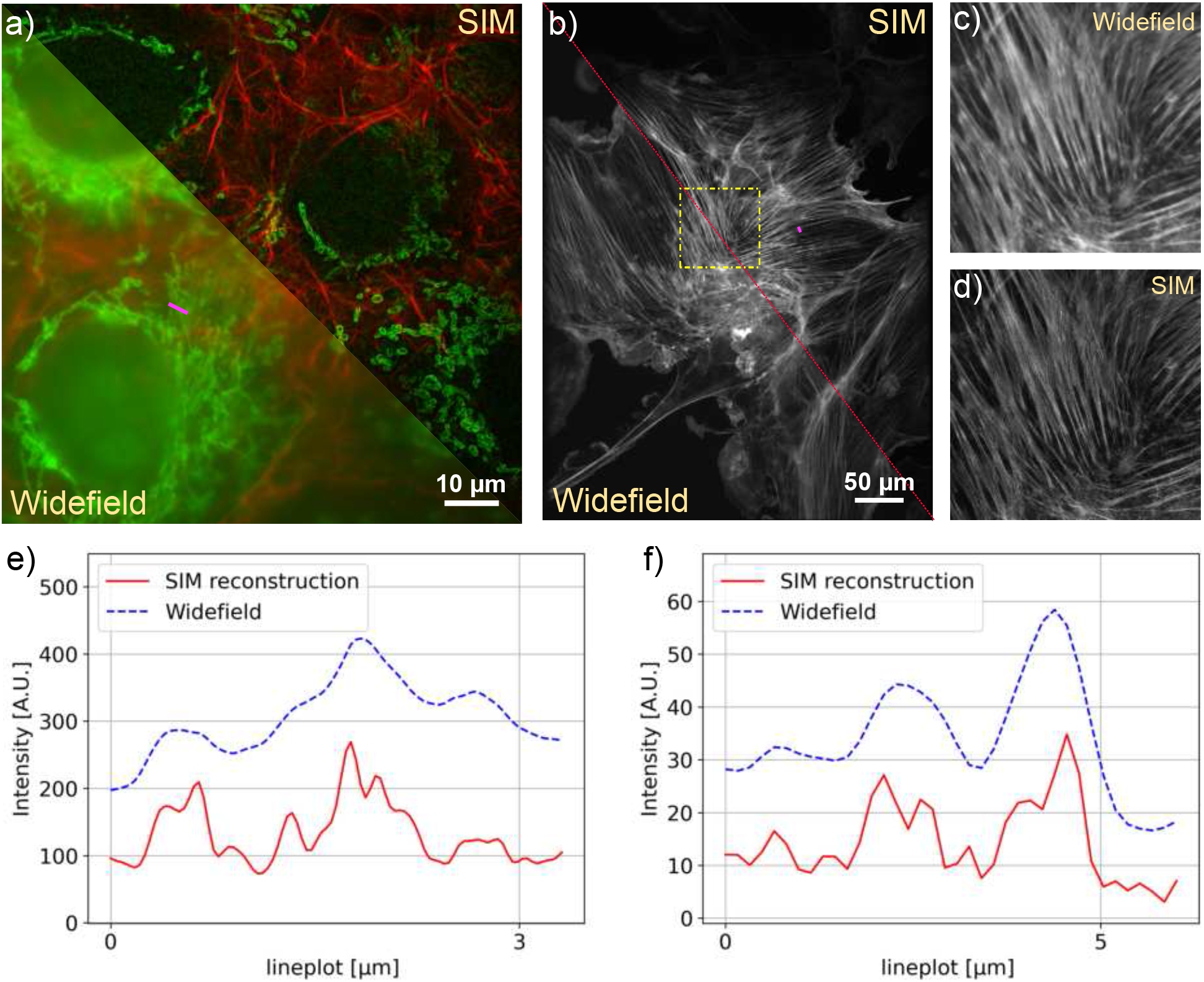
A) SIM Reconstruction and widefield comparison of Actin (488nm) and Mitochondria (635nm). The sample is GATTA-Cells 4C with stained huFIB cells (F36924, ThermoFisher Scientific, Germany). The SIM reconstruction significantly reduces the background and improves the resolution depicted in the line profile E. B) The comparison shows how SIM recovers the filamentous structure of actin in AF488 labeled samples zoomed in C/D. The sample is FluoCells prepared slide with stained BPAE cells (F36924, ThermoFisher Scientific, Germany). E) Line plot showing the widefield image and SIM reconstruction of mitochondria from A along the magenta line. F) Line plot showing the actin filaments from B along the magenta line. The SIM reconstruction separates previously unresolved filament structures.

## Appendix S7 Hardware and Software integration using ImSwitch

**FIG. S5.**
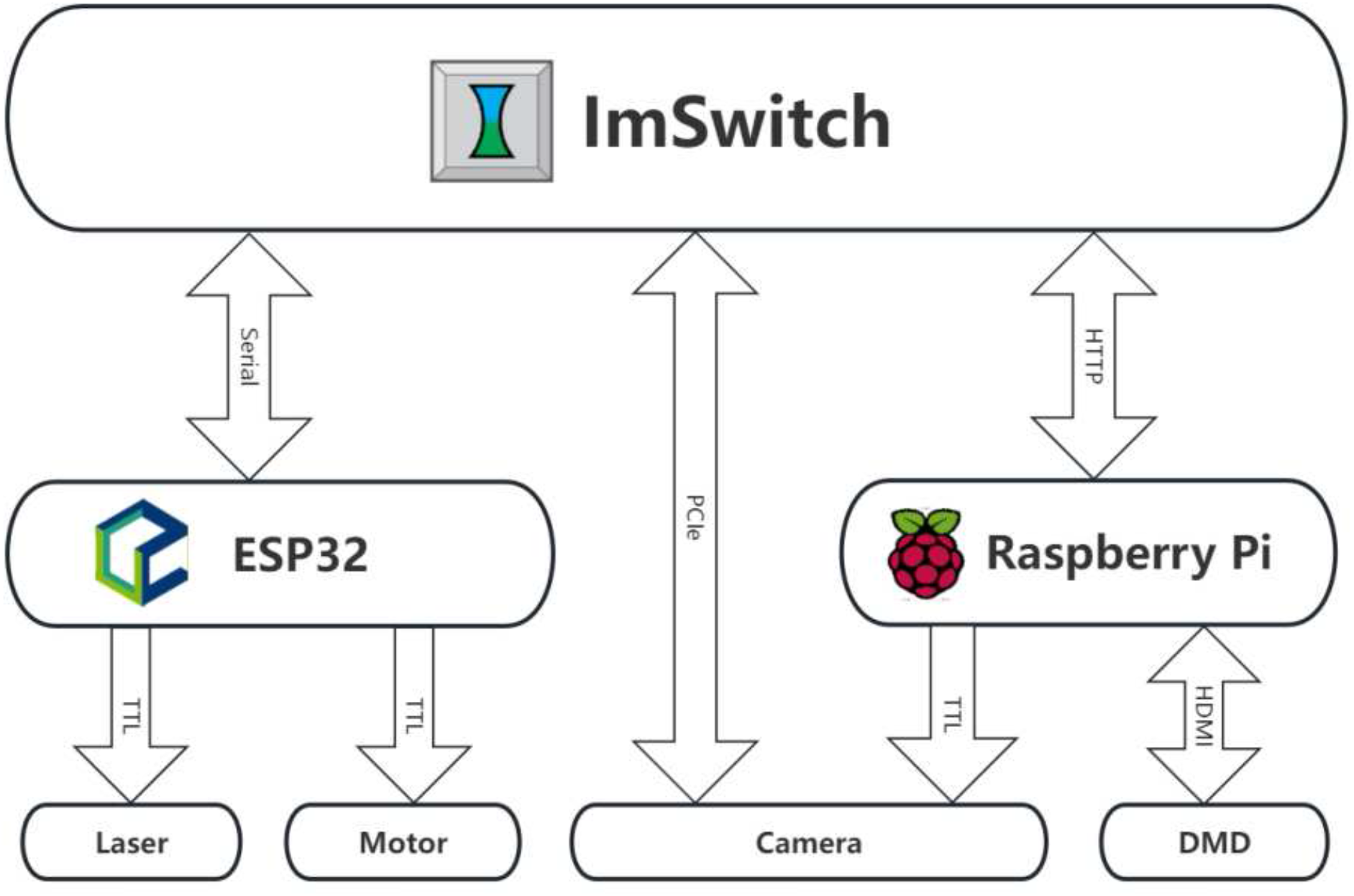
ImSwitch is the hardware orchestration, frame acquisition and image processing software that organizes displaying of the SIM patterns, motor control, camera frame readout and real-time SIM image reconstruction. Device orchestration is facilitated by ImSwitch, with a Raspberry Pi handling rapid pattern display on the DMD via HDMI and triggering the camera, while UC2-REST manages additional hardware controls, such as syncing focus position and laser intensity.

## Appendix S8 Bill of Materials

An up-to-date version of this list can be found in the online documentation https://opensimmo.github.io/docs/02_1_BillOfMaterials. Design and CAD files required for 3D printing and laser cutting custom parts are available at https://opensimmo.github.io/docs/02_2_Preparation. This also involves the creation of the Fourier masks (https://opensimmo.github.io/docs/02_2_Preparation#fourier-mask.

**TABLE S3.** Bill of Materials. Abbreviations in description indicate where part appears in the system diagram, Fig. 1.

| ID | Part | Description | Qty | Cost (€) | Link |
| --- | --- | --- | --- | --- | --- |
|  | Acrylic glass | 6mm thickness, 60x60mm, black for enclosure | 3 | 50 | Kunststoffplattenonline |
| DLP4710EVM-G2 | DMD | Pattern generation | 1 | 1100 | Texas Instruments |
| AC254-050-A | Achromatic lens | Laser collimation (L1/L2) | 2 | 80 | Thorlabs |
| AC254-075-A | Achromatic lens | Telescope (L3/L4) | 2 | 80 | Thorlabs |
| AC254-200-A | Achromatic lens | Tube lens | 1 | 80 | Thorlabs |
| PF10-03-P01 | Silver coated mirror | Laser direction (M1-M3) | 3 | 50 | Thorlabs |
| KM100 | Kinematic mirror mount | M1-M3 | 4 | 38.04 | Thorlabs |
| FL488-30 | Fiber coupled diode laser | 488nm 30mW, 4 μm core | 1 | 250 | openUC2 |
| FL635-60 | Fiber coupled diode laser | 635nm 60mW, 4 μm core | 1 | 250 | openUC2 |
| CP33/M | Optomechanical cage plate | Optical parts mount | 6 | 17.35 | Thorlabs |
| SM1FC | Fiber adapter | Fiber adapter (LD1/LD2) | 2 | 30.7 | Thorlabs |
| CXY1A | XY translation mount | Positioning Fourier mask | 1 | 183.14 | Thorlabs |
|  | Fourier mask | Aluminum foil pierced with pin | 1 |  | custom |
| SM1L40 | Lens Tube | Tube lens mount | 1 | 45.12 | Thorlabs |
|  | Raspberry Pi 3B+ | 4x 1,4 GHz, 1 GB RAM | 1 | 38.65 | Reichelt |
| ESP32-D1-R32 | ESP32 | Compatible with CNC controllerboard | 1 | 9.99 | Amazon |
| A4988 | CNC-Controllerboard | Motor driver | 1 | 17.7 | Reichelt |
| 17HS4417P1X4 | Nema17 Motor | z direction control | 1 | 15 | Reichelt |
|  | GT2 150mm | Focus control belt | 1 | 2.40 | I3DService |
|  | Pulley GT2 | Stepper motor pulley | 1 | 1.68 | I3DService |
|  | HDMI Cable | Display connection | 1 | 2.7 | Reichelt |
| RPI-PS-12.5EU-WT | microUSB powersupply | 5.1V/2.5A for Raspberry pi | 1 | 8.4 | Reichelt |
| DIN912 | M3 Screw Set | Mechanical parts montage | 1 | 15 | Amazon |
| P200/M | Mounting Post | Enclosure height | 3 | 60.86 | Thorlabs |
| PF125B | Clamping Fork | Post fixation | 3 | 14.68 | Thorlabs |
| C1515/M | Mounting Post Bracket | Enclosure leviation | 3 | 106.91 | Thorlabs |
| DIN EN 573-3 | Aluminium Profile | 20mm mount for RailOptics | 3 | 7 | Profilzuschnitt24 |

**FIG. S6.**
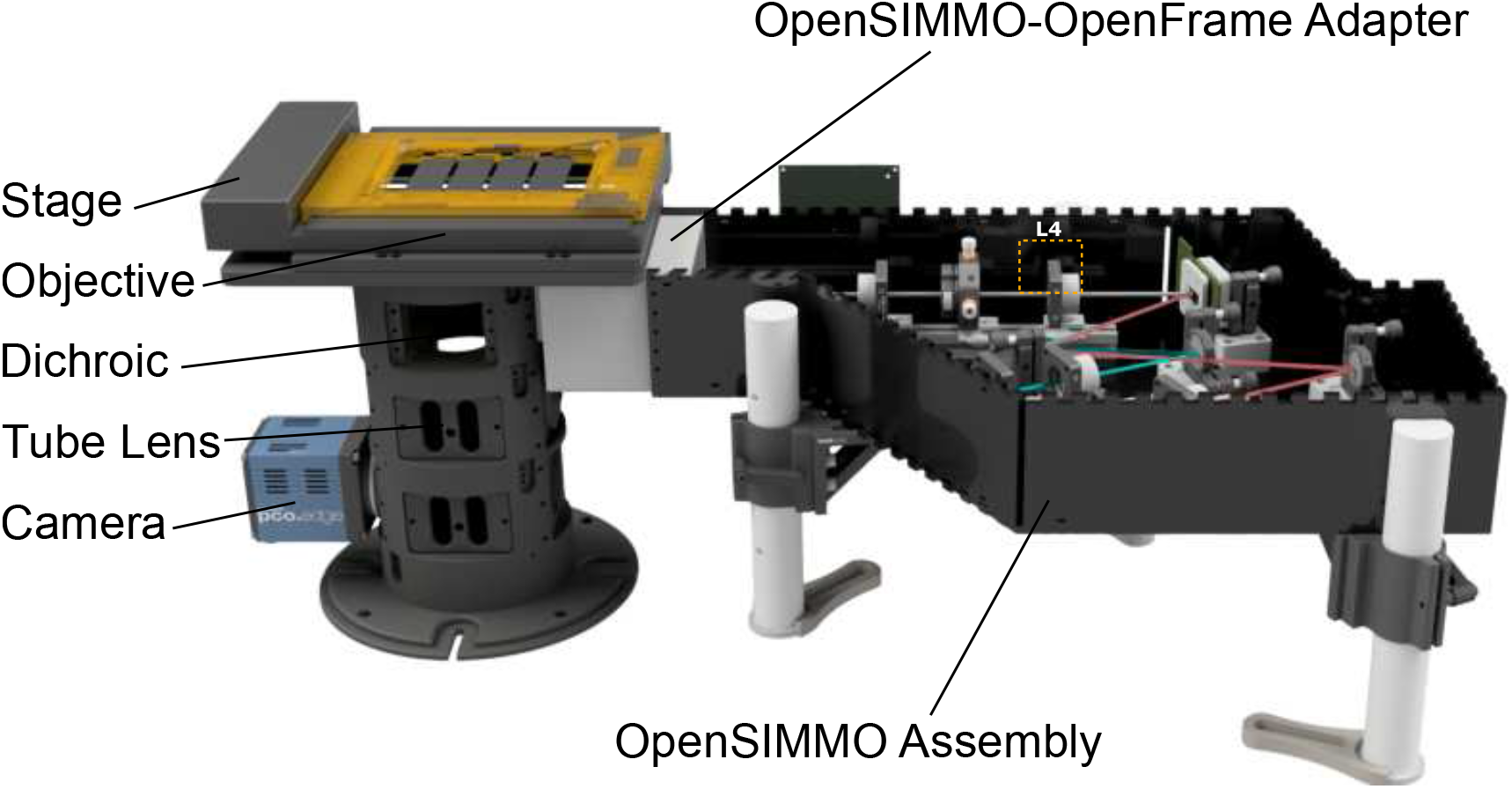
The openSIMMO setup can easily be adapted to other microscope bodies. By teaming up with other open-source projects like OpenFrame, one can build a powerful super-resolution microscope for a modest budget, with full control over the hardware and a reasonable amount of effort to build it. The expertise needed to replicate such a setup no longer requires an engineering degree. The support of the scientific community can help solve problems and improve the design over time. The modular design of the OpenFrame enables the integration of custom modules for e.g. autofocus or other fluorescent techniques.

